# Ultrastructural analysis of dendritic spine necks reveals a continuum of spine morphologies

**DOI:** 10.1101/2021.02.18.431725

**Authors:** Netanel Ofer, Daniel R. Berger, Narayanan Kasthuri, Jeff W. Lichtman, Rafael Yuste

**Author notes:** **Correspondence** Netanel Ofer, 902 NWC Building, 550, West 120 Street, Box 4822, New York, NY, 10027.

## Abstract

Dendritic spines are membranous protrusions, with a bulbous head connected to the dendrite by a thin neck, and receive essentially all excitatory inputs in most mammalian neurons. Spines have a wide variety of morphologies that likely have a significant effect on their biochemical and electrical properties. The question of whether spines belong to distinct morphological or functional subtypes or constitute a continuum is still open. To discern this, it is important to measure spine necks objectively. Recent advances in electron microscopy enable automatic reconstructions of 3D spines with nanometer precision. Analyzing ultrastructural reconstructions from mouse neocortical neurons with computer vision algorithms, we demonstrate that the vast majority of spines can be rigorously separated into head and neck components. Analysis of the head and neck morphologies reveals a continuous distribution of parameters. The spine neck diameter, but not the neck length, was correlated with the head volume. Spines with larger head volumes often had a spine apparatus and pairs of spines in a post-synaptic cell contacted by the same axon had similar head volumes. Our data are consistent with a lack of morphological categories of spines and indicate that the morphologies of the spine neck and head are independently regulated. These results have repercussions for our understanding of the function of dendritic spines in neuronal circuits.

## 1 INTRODUCTION

Dendritic spines, small neuronal appendages that mediate essentially all excitatory transmission in the brain, were discovered by Cajal more than a century ago (Cajal, 1904), who noted the existence of a ‘ball-like’ spine head and a ‘lightly stained’ spine neck (Yuste, 2010). However, these findings were controversial and not proven until the inventions of the electron microscopy (EM) that allows imaging of dendritic spines with nanoscale resolution (Gray, 1959). Recent advances in EM allow automatic imaging of every neuron in a brain tissue volume, including high-resolution 3D structure of spines (Kasthuri et al., 2015; Dorkenwald et al., 2019; Lee et al., 2019; Motta et al., 2019). The 3D ultrastructure is important since spines, and the border between the head and neck, appear different from different angles of observation (Arellano et al., 2007). The question of whether spines belong to different classes or are part of a continuum is still controversial (Pchitskaya and Bezprozvanny, 2020), and part of the problem lies on the lack of a clear definition of what constitutes the spine neck. Peters and Kaiserman-Abramof examined the spines of rat neocortex, proposing the distinction between stubby, mushroom, and thin spines (Peters and Kaiserman-Abramof, 1970). They defined stubby spines as short thick spines without a clear neck, mushroom spines as thick necks that expands into a large end-bulb, and thin spines as slender necks that expands into a small oval or rounded end-bulb. This classification has been widely adopted and used in almost every study on spines at the optical microscope level (Arellano et al., 2007). However, no clear borderline was observed between these three types and a continuum and unimodal distribution of the spine’s morphological parameters has been reported in several studies (Peters and Kaiserman-Abramof, 1970; Arellano et al., 2007; Tønnesen et al., 2014; Loewenstein et al., 2015). Recently, analyses of large-scale connectomics datasets of mouse cortex quantified synaptic size (Motta et al., 2019) and spine head volume (Dorkenwald et al., 2019) and described two subtypes of spines: small and large. However, there was significant overlap in sizes between them. Thus, the question of whether dendritic spines are divided into distinct types or constitutes a continuum is still undecided (Rochefort and Konnerth, 2012; Berry and Nedivi, 2017). This controversy is confounded by the lack of methods to define and measure spine necks objectively.

In this study, we analyzed an ultrastructural dataset from mouse neocortex and introduce computer vision algorithms to show that the vast majority of spines present a clear morphological separation between head and neck. Then, we used these morphological parameters to examine whether distinct types of spines can be detected, finding unimodal and continuous distribution of spine parameters. Finally, we analyzed the correlation between spine morphological properties, including the post-synaptic densities (PSD) and spine apparatus (SA), and find a lack of correlation between the spine neck length and the spine head, indicating that the two compartments are independently regulated. Our methods and results can serve to further explore the development and function of spines.

## 2 METHODS

### 2.1 Ultrastructural Dataset

In this paper, we analyzed a set of 4,223 dendritic spines of layer 6 pyramidal cells (at the apical dendrite level in layer 5) from the somatosensory cortex of a young adult mouse, published in (Kasthuri et al., 2015). The EM data was acquired using automated ultrastructural technology with nanometer resolution. The detailed methodology to reconstruct spines in three dimensions is described in Kasthuri et al. (2015). Neuronal processes in this image dataset were segmented using VAST, a computer-assisted manual space-filling segmentation and annotation program (Berger et al., 2018). VAST allows coloring images at multiple scales of resolution, to organize the results in a flexible annotation framework, and to export results for 3D visualization and analysis. Each spine was represented by a triangle mesh in an ‘OBJ’ format which includes a list of 3D vertices followed by a list of faces formed by the vertices (Figure 1A). A detailed spreadsheet for 1,700 synapses, that describes synapse position, axon ID, dendrite ID, and biological details, such as if the spine consists an SA and PSD size, is provided by Kasthuri et al. (2015). The IDs of the pre- and post-synaptic partners enabled us to find pairs of spines that create dual connection between the same pre- and post-synaptic neurons.

**FIGURE 1.**
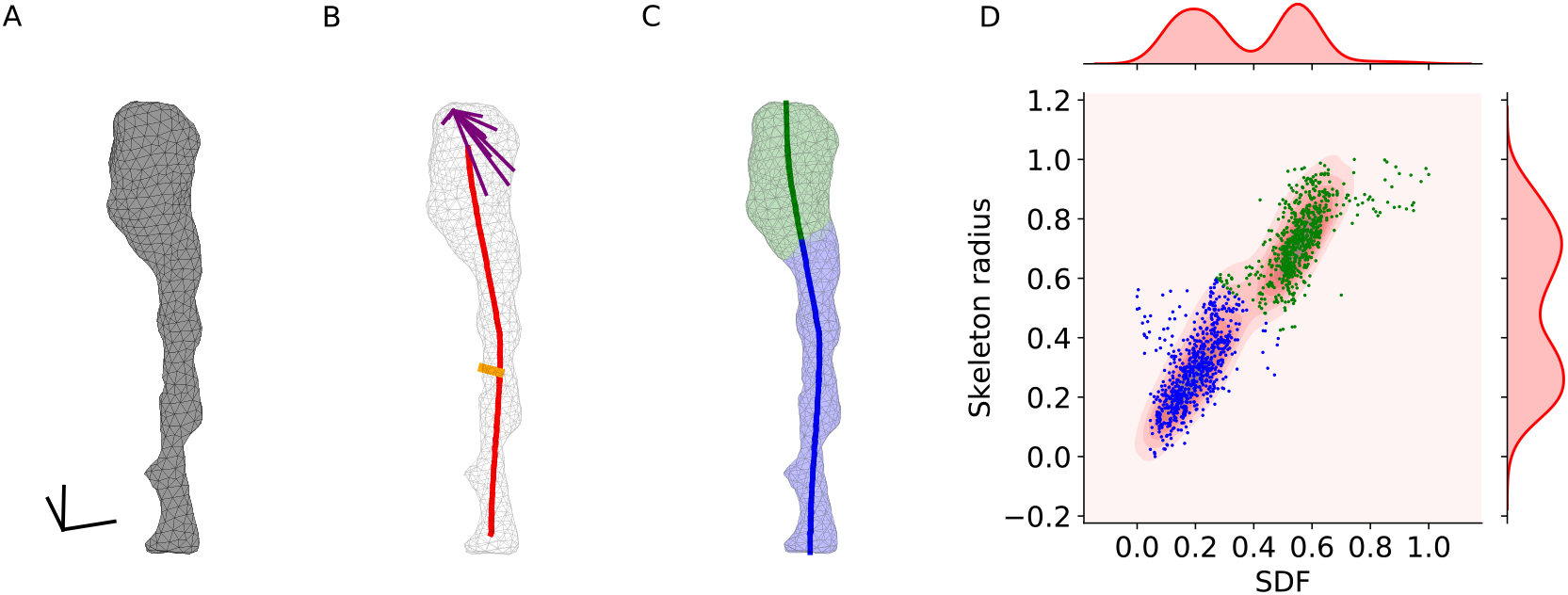
Computational separation of spine head and neck. (A) Triangle mesh of a spine, scale bar: 100*nm*. (B) Centerline skeleton curve is colored in red. An example of the ’skeleton radius’ is colored in orange. An example of rays in a 60° cone for measuring the SDF colored in purple. (C) The head (green) and neck (blue) were classified according to the SDF and ’skeleton radius’ values. The centerline curve was extended and divided into neck length (blue) and head length (green). Parameters values are: spine volume: 0.006*μm*^3^, head volume: 0.004*μm*^3^, spine surface area: 0.268*μm*^2^, head surface area: 0.141*μm*^2^, spine length: 0.854*μm*, neck length: 0.596*μm*, neck radius: 0.036*μm*, head sphericity: 0.864. (D) Scatter plot of faces SDF and ’skeleton radius’ values. Each dot represents a single face of the spine. Distributions of SDF and ’skeleton radius’ on top and right. Faces were clustered using Gaussian Mixture Model. Hartigan’s dip-test p-values are under 0.001, indicating bimodality. The spine ID: Kasthuri_4643.

### 2.2 Morphological Analysis

In our first analysis stage, Laplacian smoothing, each vertex in a mesh covering each spine was replaced by the average of its neighbors (that share an edge). This was applied to compensate for the quantization effect caused by the EM sectioning process. Next, small isolated components that contain less than 17 faces were removed. Shape Diameter Function (SDF) and Mesh Skeletons were then calculated using the ‘Triangulated Surface Mesh Segmentation’ and ‘Triangulated Surface Mesh Skeletonization’ packages from the Computational Geometry Algorithms Library (CGAL) 5.0.2, https://www.cgal.org (Shapira et al., 2008; Tagliasacchi et al., 2012). To this end, data were converted from ‘OBJ’ to ‘OFF’ format. SDF measurement were done by averaging 25 rays beams in a cone of 60 degrees projecting to the opposite spine surface (purple rays in Figure 1B). The SDF is a pose-independent method that matters for this dataset because of the tortuous nature of spines. The ‘skeleton radius’ is the distance from each face to the closest point along the mesh skeleton (orange line in Figure 1B). These algorithms, normalized between 0 and 1, were then used for the segmentation between the head and neck, and later, for measurements of the spine morphological parameters.

For the segmentation of spines into head and neck, we used two local parameters for each face, the SDF and the ‘skeleton radius’ values, as well as a spatial parameter, the dihedral angle between neighboring faces. The energy-function algorithm applied a graph-cut-based algorithm that combines fast changes on SDF and ‘skeleton radius’ values as natural candidates for segment boundaries and geometric criterion in adjacent facets, sharing a sharp and concave edge (CGAL ‘Triangulated Surface Mesh Segmentation’ package). The parameters used in this algorithm were the number of clusters of 2 and smoothing-lambda of 0.1.

### 2.3 Morphological Parameter Measurements

The volume of spines was calculated using the signed tetrahedral volumes summarizing algorithm (Zhang and Chen, 2001):

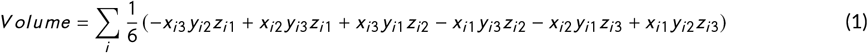

where (*x*_*i* 1_, *y*_*i* 1_, *z*_*i* 1_), (*x*_*i* 2_, *y*_*i* 2_, *z*_*i* 2_), and (*x*_*i* 3_, *y*_*i* 3_, *z*_*i* 3_) are the coordinates of the vertices of face *i*. This algorithm requires a closed mesh and face normals pointed to the same inner or outer direction (the order of the face vertices, clockwise or counterclockwise, indicates the normal). Thus, for calculating the spine head volume, after computationally severing the head, the hole was filled by connecting the border faces to the center point of the hole, considering the neighbor faces to find the correct direction of the normals.

For measuring spine length, we summed the length of the skeleton center line and the extension to the spine surface, in the direction of the vector of the last two vertices, in the two edges. For measuring spine’s neck length separately, each vertex among the centerline was labeled as ‘head’ or ‘neck’ according to the major of its belonged vertices. Summarizing the ‘neck’ segments gives us the neck length.

The spine head sphericity was calculated by equation (2):

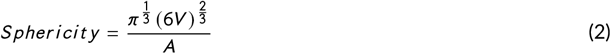

where V is the spine head volume and A is the area. In a perfect ball, the sphericity equals 1.

### 2.4 Statistical analysis

Kolmogorov-Smirnov test for two samples was used to compare two empirical cumulative distribution function (CDF) with a two-tailed p-value. The Hartigan’s dip-test of unimodality (Hartigan and Hartigan, 1985) was applied to examine whether data are unimodal distributed. Since the Hartigan test was designed for a 1-dimensional dataset, for considering mutually two parameters, we tested the unimodality on 18 projections of 10-degree rotations of the 2-dimensional dataset (Schelling and Plant, 2020).

### 2.5 Code accessibility

The CGAL scripts were written in C++; the other codes were written in Python 3.7 using the libraries: numpy 1.17.4, scipy 1.5.4, scikit-learn 0.23.1, and unidip 0.1.1. All the data is publicly available at the Columbia University Academic Commons site (https://academiccommons.columbia.edu). Codes used in this paper are publicly available at the Columbia University Neurotechnology Center’s GitHub page (https://github.com/NTCColumbia).

## 3 RESULTS

### 3.1 Computational separation of spine head and neck

We first explored the morphological identity of the spine neck. To do so, we segmented spine structures into head and neck using their morphological properties. To this end, we divided the surface of the spine into a triangle mesh, and for each face of the spine surface, two geometrical parameters, the SDF and the ‘skeleton radius’, were calculated. The SDF and the ‘skeleton radius’ are complementary morphological parameters, and the combination of both distributions enabled a robust segmentation between the head and neck, as evident visually in their bimodal distribution of values (Figure 1D). The cluster with the lower value of average SDF was labeled as ‘neck’ (blue) and the other cluster was labeled as ‘head’ (green). To quantify the extent of the separation between head and neck for every spine, we tested whether the distributions of the SDF and the ‘skeleton radius’ values were unimodal. Denying the possibility of unimodality implies a bimodal (or higher order) distribution, with a clear clustering into two groups: the head and the neck faces. Statistical tests were applied to examine if the separation between head and neck was significant. The Hartigan’s dip-test revealed that 88.44% (3,717/4,203) of spines had a bimodal distribution of SDFs, and 74.87% (3,147/4,203) of spines had a bimodal distribution of ‘skeleton radius’ values (p-value < 0.05, Hartigan’s dip-test). Mutual SDF and ‘skeleton radius’ 2-dimensional Hartigan’s dip-test (see Methods) led to 95.12% (3,998/4,203) of the spines demonstrating statistically significant bimodal distributions. To find the exact border between head and neck we used a graph-cut algorithm with energy function minimization, taking into account the SDF and ‘skeleton radius’ values as well as the convexity of the mesh surface. The border was often in sharp dihedral angles between neighboring faces, which occur between the head and neck. In 74.33% of the spines, the two clusters of faces led to two segments, one for head faces and the other for the neck faces. In 1% of spines there was only a single cluster, meaning that there was no separation between head and neck. In the rest of the spines, the two clusters led to more than two segments. These cases included branched spines in which two connected thin cylinders branched into two different heads and also cases of two mistakably coupled spines. We concluded that most spines can be rigorously separated into a head and neck. For the rest of the study, focused on the morphological analysis of spine necks, we analyzed only spines consisted of clear two segments (3,138/4,223 spines).

### 3.2 A continuum of spines morphological parameters

We then proceed to build a dataset of different morphological variables for each spine. First, we measured three basic morphological parameters of the entire spine: its volume, surface area, and length. The spine surface area was measured by summarizing all the triangle mesh areas of the spine. The volume of the spine was calculated using the signed tetrahedral volumes algorithm (Zhang and Chen, 2001). The spine’s length was obtained by measuring the extended centerline skeleton straightforward on the two sides (Figure 1C). The separation between the spine head and neck enabled us to also measure the ‘head volume’, ‘neck length’, and ‘neck radius’. The head surface area was measured by integrating the area of all faces labeled as a ‘head’. Since the algorithm for volume measurements requires a closed volume, we first filled the hole created by the cutting of the neck with a simple plane, followed by calculating the volume of the new closed spine head mesh. For accurate measurement of neck length, we used similar methods as with total spine lengths. Each vertex along the skeleton centerline was labeled as ‘head’ or ‘neck’ according to most of its faces (Figure 1C). Thus, we integrated the lengths along the neck-labeled vertices along the centerline curve. In contrast to previous methods, that measure Euclidean or Geodesic distances, either manually or semi-automatically with user mediation (Jorstad et al., 2014, 2018), our method is a unique automatic approach for accurate measuring of the spine neck, because it measures the neck length along the center of the 3D spine, considering also the head position. Finally, to measure neck radius, we averaged the shortest distance between the face center and the centerline skeleton curve for all of the faces belonging to each vertex along the skeleton centerline (see orange line in Figure 1B, as example of a neck radius for a specific face). Then, we averaged the averaged values along all the vertices. This method for measuring spine neck dimensions provided the most accurate results, even for non-round (elliptic) neck cross-sections. Neck radius was perpendicular to the centerline skeleton, so it was not affected by the 2D section-cutting or projection angle. The measured values of the morphological parameters are summarized in Figure 2A-G. Inspection of the distributions for each of the morphological parameters revealed skewed unimodal functions, with no clear bi- or multimodality. To explore this in depth, and test if spines could be classified into different morphological subtypes, we plotted the data along pairs of variables, including ‘spine volume’, ‘spine surface area’, and ‘spine length’ (Figure 3A), and ‘head volume’, ‘neck length’, and ‘neck radius’ (Figure 3B). Visual inspection failed to show clear multimodal distributions. This was confirmed with the 3-dimensional Hartigan’s dip-test, finding a continuous and unimodal distribution. With this lack of multimodality in the morphological parameters, we cannot prove the existence of distinct spine types and concluded that, in our dataset, spines displayed a continuum distribution of morphologies.

**FIGURE 2.**
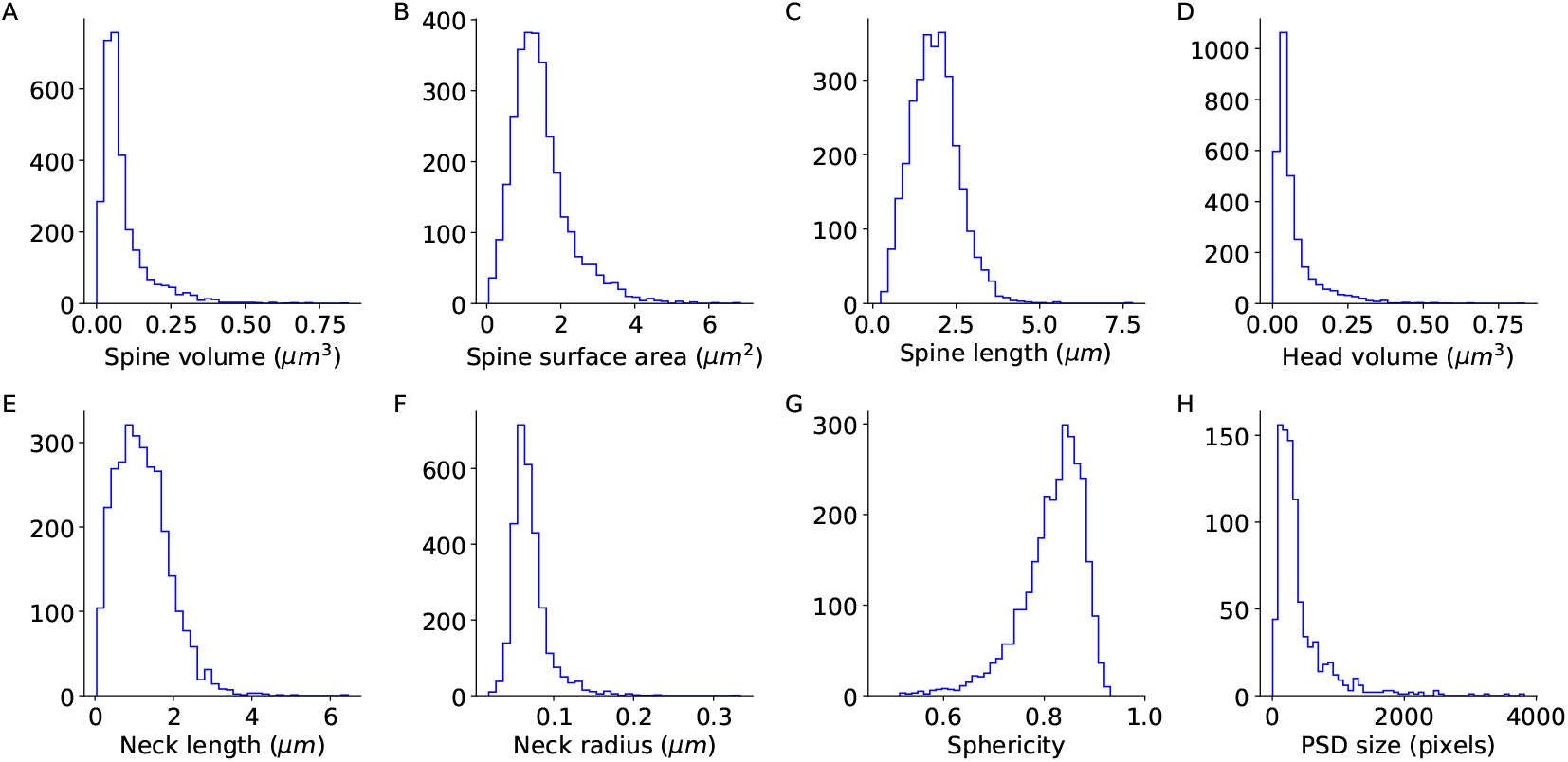
Unimodality of spine morphological parameters. (A-C) The morphological parameters of the entire spine. (D-F) The morphological parameters for the separated head and neck. (G) The sphericity of the head volumes. The dataset includes 2,998 spines. (H) Post-synaptic density size distribution, a dataset of 888 spines.

**FIGURE 3.**
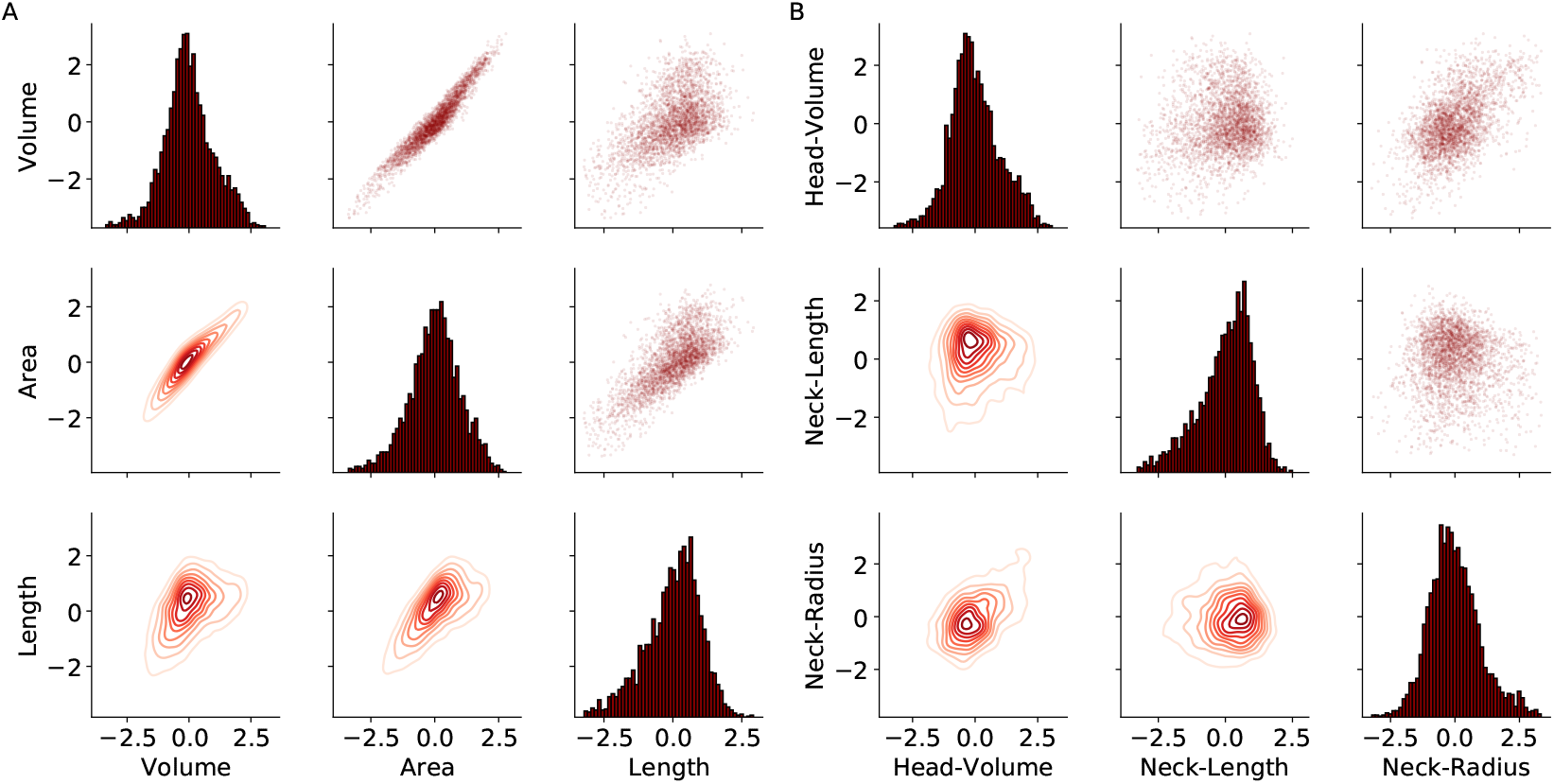
Continuum distribution of spine morphological parameters. The pair-plot presents in the upper and lower triangles the pairwise relationships of the morphological parameters of all the spines. The marginal distribution of each parameter can be shown on the diagonal. (A) For the three basic parameters of the entire spine. (B) For neck and head parameters after segmentation. The data were logarithmic z-scored and the outliers (above 3 STD) were removed.

### 3.3 Correlations between spine morphological parameters

The morphological ratio between head and neck affects the electrical and biochemical isolation of the spine (Araya et al., 2006). Previous studies have reported no correlation between spine head and neck parameters (Araya et al., 2014; Tønnesen et al., 2014). Other studies found a weak correlation between head volume and neck diameter in chemically fixed spines, but not in cryo-fixed spines (Arellano et al., 2007; Tamada et al., 2020). We used our dataset and analysis pipeline to explore this issue with a larger number of ultrastructural reconstructed spines (Figure 4).

**FIGURE 4.**
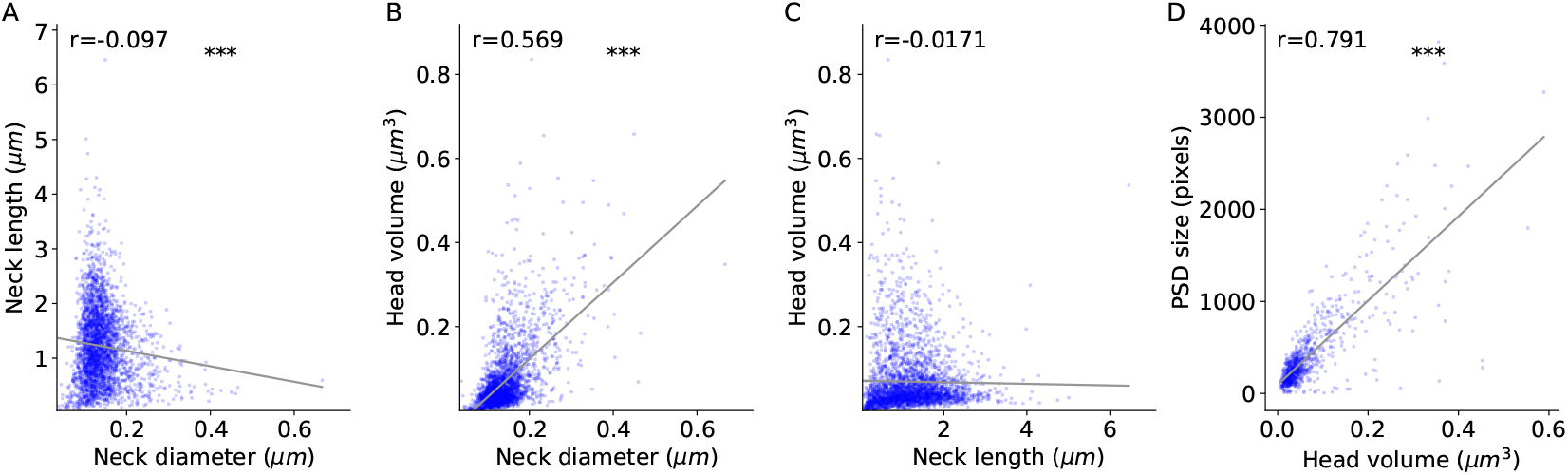
Correlation between spine head and neck morphological variables. (A-C) Correlation between head and neck morphologies (2,998 spines). (D) Correlation between head volume and post-synaptic density size (888 spines). The correlation coefficients (Spearman) are indicated for each graph. The asterisks indicate statistical significance *** *p* < 0.001. Two-sided p-value for a hypothesis test whose null hypothesis is that the slope is zero, using Wald Test with t-distribution of the test statistic.

Our results confirmed the existence of a strong correlation between head volume and neck diameter (Figure 4B), a weak negative correlation between neck diameter and neck length (Figure 4A), and a lack of a significant correlation between head volume and neck length (Figure 4C). In addition, to investigate the relationship between biological properties and physical dimensions, we analyzed the PSD size and presence or absence of an SA, as a function of the measured spine morphology. In agreement with previous studies (Arellano et al., 2007; Holler et al., 2021), the PSD size presents a long-tailed unimodal distribution (Figure 2H), and was strongly correlated with the spine head volume (Figure 4D). We then examined the relationship between the presence of an SA and spine morphology. Spines with SA had higher head volumes and neck diameters than spines without SA (Figure 5). This was particularly apparent in cumulative distribution functions (Figure 5B), which showed also no differences between spines with SA and spine without SA in neck length (Figure 4C).

**FIGURE 5.**
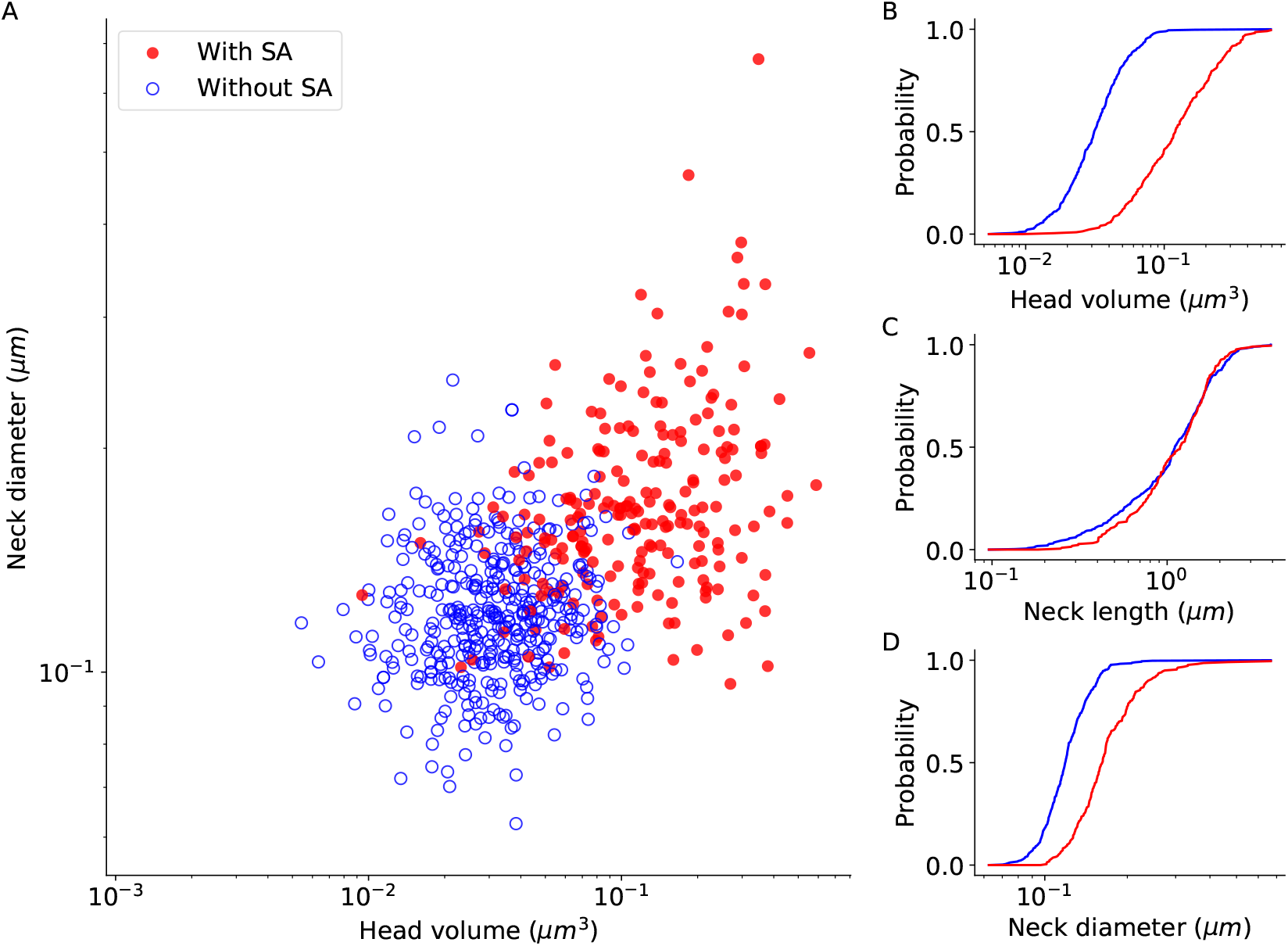
Spine apparatus is present in larger spines. (A) Spine apparatus as a function of head volume and neck diameter. Full red circles indicate the spine with spine apparatus, and empty blue circles indicate the spine without spine apparatus. (B-D) The empirical cumulative distribution function of the spines with spine apparatus (red) and without spine apparatus (blue). Kolmogorov-Smirnov test p-values: (B) *p* < 0.001, (C) *p* = 0.3677, and (D) *p* < 0.001. Spine apparatus indicated as ‘uncertain’ or ‘N/A’ in the spreadsheet were not included, resulted in 401 spines with spine apparatus, and 220 spines without spine apparatus.

Finally, we examined the morphologies of spines that create dual connections, sharing the same pre- and post-synaptic neurons, by calculating the difference and the ratio of the morphological parameters between dual connection spines. We compared these distributions to the difference and ratio of any two spines, randomly chosen from the entire dataset. To examine whether the underlying probability distributions of the two empirical CDF curves differ, the Kolmogorov-Smirnov test was used. The differences between the morphological parameter values showed the same distribution as for two random spines (Figure 6A-C). For neck length and neck diameter, the ratio between the dual connection spines also resembled those from two random spines (Figure 6E-F). However, the ratio between head volumes in dual connection spines were lower than those of two random spines (Figure 6D). These results are in line with previous studies that reported a correlation in head volumes between the two spines of a dual connection (Kasthuri et al., 2015; Dorkenwald et al., 2019; Motta et al., 2019).

**FIGURE 6.**
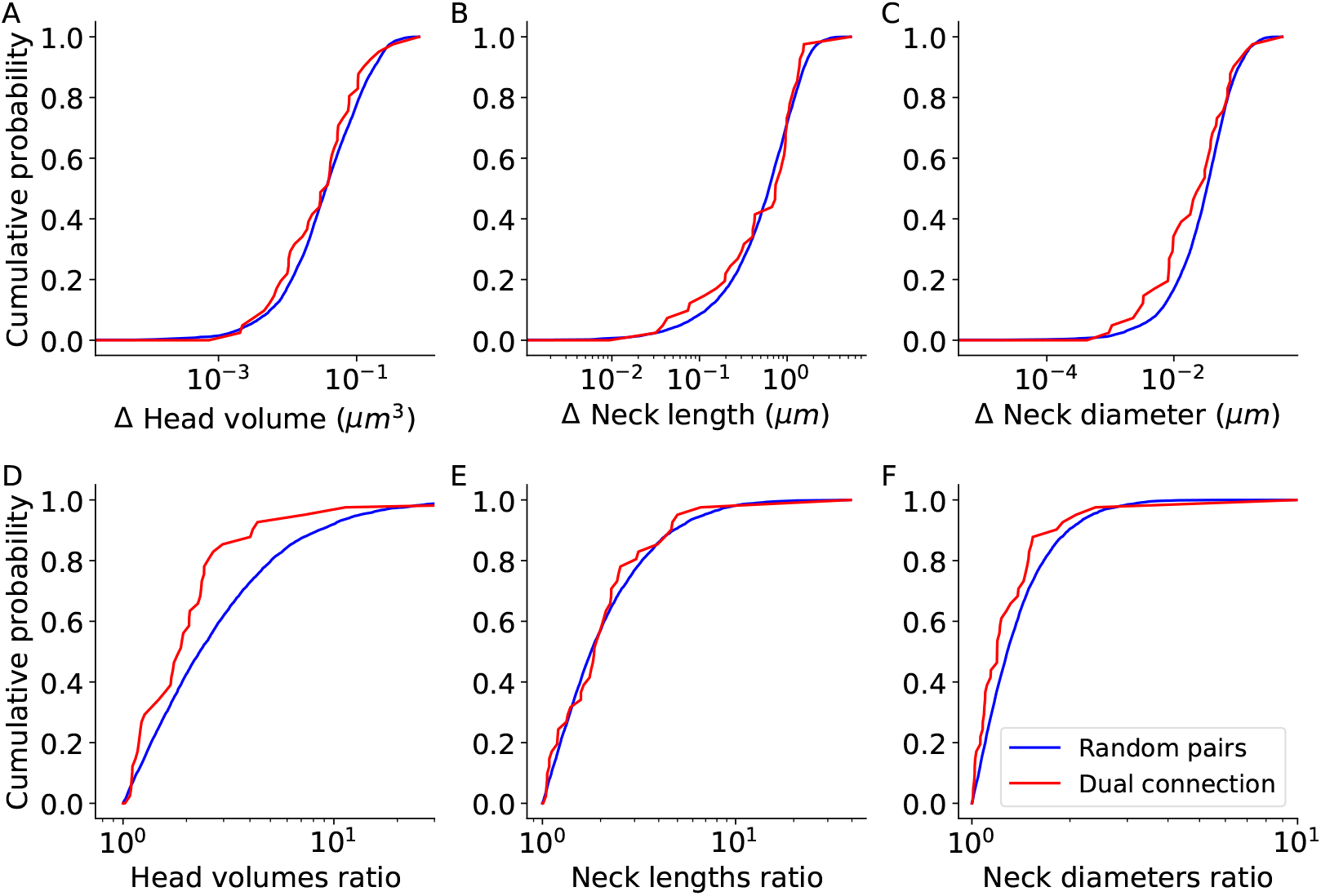
Dual connection spines have similar head volumes. The empirical cumulative distribution function of the difference between two spines (A-C) and the ratio between two spines (D-F) belong to the dual connection (red), compared to two random spines from the entire database (blue). Kolmogorov-Smirnov test p-values: (A) *p* = 0.6475, (B) *p* = 0.7694, (C) *p* = 0.0671, (D) *p* = 0.003, (E) *p* = 0.5915, and (F) *p* = 0.1046. The dataset includes 41 dual connection pairs.

## 4 DISCUSSION

Dendritic spines display a large morphological heterogeneity of head and neck dimensions. These differences likely have functional meaning, as the neck can cause a biochemical and electrical isolation between the spine head and the dendritic shaft. Since spines regulate most excitatory communication between neurons, this diversity could enrich the computational capabilities of the brain. Here, using an objective head and neck separation algorithm, we showed that the vast majority of spines have a clear head and neck. Based on this, we developed new methods to measure the spine head volume, neck length, and neck diameter. These morphological parameters present a continuum distribution in our dataset, in agreement with previous proposals that spines do not belong to different morphological subtypes. Finally, we examined the correlation between the morphological parameters and the relationship between them and the PSD size and SA and detect correlations between spine volume and neck width, PSD size and presence of SA, but find no correlation between neck length and head volume.

### 4.1 Objective identification of spine necks

Although the spine’s head and neck dimensions determine its electrical and chemical activity (Segev and Rall, 1998; Yuste, 2013; Bell et al., 2019; Lagache et al., 2019), the definition of head and neck and the exact border between them has always been unclear. Previous studies measured the head and neck manually or using algorithms with arbitrary cutoffs that required a large number of corrections by a human annotator (Benavides-Piccione et al., 2002; Arellano et al., 2007; Dorkenwald et al., 2019; Motta et al., 2019). In this study, we developed an automatic algorithm to separate between head and neck, confirming that spine neck is real and that, in the vast majority of spines, one can separate the head and neck in a statistically significant manner.

### 4.2 A continuum of spine morphologies

Given the great variety of spine morphologies, an open question in the field is whether spines belong to different morphological subtypes. The common nomenclature of Peters and Kaiserman-Abramof classified spines into three types, stubby, mushroom, and thin (Peters and Kaiserman-Abramof, 1970). Their description of stubby spines, without a well-defined neck, may be wrongly reported due to limited spatial resolution of optical microscopy (Tønnesen et al., 2014). In fact, ‘stubby’ spines without a clear neck are very rare, approximately 1% in our dataset. Their ‘thin’ spines were originally named after their ‘slender stalk’, without taking into account the variety of neck lengths, so one could question the validity of that term. Following Peters and Kaiserman-Abramof visual classification, semi-supervised learning and a decision tree have also been used to classify spines into these same types: stubby, mushroom, and thin (Rodriguez et al., 2008; Janoos et al., 2009; Shi et al., 2014; Basu et al., 2018). On the other hand, unsupervised morphology-based clustering of dendritic spines from human cortical pyramidal neurons uncovered at least six separate groups of spines (Luengo-Sanchez et al., 2018).

Our analysis revealed a clear continuum distribution of the morphological parameters, without any evidence of separate subtypes (Figure 3). Statistical tests applied to these data, the same that proved the morphological reality of spine neck and heads, do not reject the unimodal hypothesis, meaning that we cannot prove the existence of distinct types of spines. This conclusion is of course limited to our dataset and our measured variables, so we cannot rule out the possibility that in different datasets, or with different morphological measurements, one could identify different subtypes of spines. Also, different morphological subtypes of spines could exist but have overlapping morphological parameters. For example, the boundaries between spines subtypes could be blurred as a result of the dynamic morphological transition between spine types, leading to a unimodality in cluster analysis. While theoretically possible, the simplest interpretation of our results that spines represent a continuum of morphologies, without any clear subtypes.

### 4.3 Functional considerations

Studying the morphological parameters of dendritic spines may shed light on their functional role. The lack of correlation between head volume and neck length (Figure 4C) points out the possibility of different biological mechanisms governing the development of the spine head and neck. For example, the fact that spines reach out to axons running close-by, which may dictate spine length, could explain the lack of correlation of neck length with head size. Moreover, the similarity in head volume between dual connection spines, but not in neck length (Figure 6), implies different functional roles of the head and neck during synaptic activity. The correlation between presence of SA and head volume was reported in previous studies (Dorkenwald et al., 2019). In addition, here we show also a significant relationship between SA and neck diameter (Figure 5). This could be interpreted as if the large size of the head, together with a thick neck, enables the entrance of SA from the dendritic shaft into the spine head.

Our analysis used a limited dataset of spines from pyramidal cells from the mouse somatosensory cortex (Kasthuri et al., 2015). To strength our results, and answer the questions of whether dendritic spine morphologies are continuous or group and the extent of correlation between dual connection spines, more data are needed. Future studies should expand and compare the results to other species and brain regions, particularly to examine the difference between mice and humans.

While EM reconstruction of spines have nanometer resolution, they arise from a fixed structure, probably representing snapshots of spines in morphological transition that can bias us towards a misleadingly static view of spine morphology. For studying the dynamic properties of the spine and tracking structure changes, the usage of superresolution microscopy data should be considered.

In closing, the rich diversity of spine morphologies must enrich the neuronal circuit function. The computational advantages of adding such complexity should be studied theoretically and by models. Artificial neural network with realistic architecture could be used to not only to model brain circuits but, in addition, to explore improvements of existing algorithm in a brain-inspired manner.

## Acknowledgements

Supported by the NINDS (R01NS110422; R34NS116740). This material is based upon work supported by, or in part by, the U. S. Army Research Laboratory and the U. S. Army Research Office under contract number W911NF-12-1-0594 (MURI).

## Author Contributions

N.O. and R.Y. conceived the project. N.K. and D.B. performed experiments and N.O. and R.Y. wrote the paper. N.O., N.K. and D.B. analyzed the data. J.L. discussed results and edited the paper. R.Y. and J.L. assembled and directed the team and secured funding and resources.

